# Specific basal ganglia, amygdala, and cortical neurons are associated with borderline personality disorder (BPD)

**DOI:** 10.64898/2026.09.04.749244

**Authors:** Hazal Senturk, Hadar Segal-Gavish, Jason Tucciarone, Laramie Duncan

## Abstract

Borderline personality disorder (BPD) is clinically significant and poorly understood biologically. We tested which human brain cell types preferentially use genes associated with BPD, implicating two major types of neurons: LAMP5 interneurons and D2 medium spiny neurons. These interrogable cell types provide new avenues for investigating the neurobiology of BPD.

## Introduction

Borderline personality disorder (BPD) is a clinically significant condition that is poorly understood biologically. It affects roughly 1–2% of the population, typically emerges in adolescence, and is defined by pervasive instability across emotion, self-image, and interpersonal relationships, together with marked impulsivity. Rates of self-harm, suicidal ideation, and suicide attempts are high, and there is high comorbidity with other psychiatric disorders. Although BPD is treatable and many patients improve with structured psychotherapy, symptom remission is often incomplete. Pharmacotherapy has limited evidence for core BPD symptoms and is generally reserved for comorbid psychiatric disorders or short-term crisis management. (1)Therefore, it is critical to better understand the neurobiology underlying BPD, such that therapeutics can be intentionally designed.

Given that BPD is, at best, modeled in animals with substantial uncertainty, reviews of the neurobiology of BPD focus primarily on human brain imaging findings, in which the most reproduced signal is likely emotionregulation related amygdala hyperactivity(2). While useful, MRI methods do not have adequate temporal and spatial resolution to support inferences about the functioning of specific cell types in the human brain. Thus, there is tremendous need for alternative methods to characterize the neurobiology of BPD, and unbiased genetics approaches may prove particularly useful for discovering mechanisms underlying BPD.

BPD is known to be moderately heritable (*h*^*2*^_*twin*_ =46%)(3) and to have high comorbidity and genetic correlations with other psychiatric disorders (e.g., PTSD, depression, OCD, schizophrenia; rg=0.86, 0.83, 0.50, and 0.49, respectively). Recently, the first reasonably well-powered genome-wide association study (GWAS) of BPD was completed, yielding 11 significant loci(4). This GWAS underscored the necessity of unbiased genomic approaches; candidate gene studies of BPD focused largely on serotonin related genes (e.g., *SLC6A4, TPH2*, and *MAOA*), and none of these genes were among those associated with BPD in the unbiased BPD GWAS. These genes ranked, #6,895, #4,842, and #726, of 19,033 genes tested. The top GWAS genes included long intergenic non-coding RNAs (*LINC00970, LINC01680*), transcription factors (*FOXP1* and *FOXP2*), and other genes not previously studied.

Here we extend the translational potential of this BPD GWAS by using the full summary statistics (versus only the top genome-wide significant loci) and by asking (statistically) which human brain cell types preferentially use BPD associated genes(5). We then depict these cell types in their primary locations in the human brain (according to best available knowledge(6)) and characterize these cell types in terms of their primary classes (e.g. D2 medium spiny neurons).

## Methods

Using reported previously methods(5), we analyzed GWAS summary statistics (6,043,895 genetic variations) for BPD to determine which, if any, human brain cell types preferentially use genes that are more associated with BPD, as determined using MAGMA(7), software that aggregates genetic signal from all genetic variants within a gene and quantifies phenotypic associations at a gene level. Cell types tested here are from the most comprehensive study of cell types in the human brain, which encompassed 3,369,219 nuclei sampled from 105 human brain regions and then clustered into 461 cell types(6). MAGMA gene property analysis was then used for the cellular association analysis, by implementing a regression framework to test for positive relationships between gene specificity and phenotypic associations, for each of the 461 cell types. Bonferroni correction was used to adjust for multiple testing (p<.05/461=.0001). This is a computational analysis that does not involve any participants, and uses only publicly available data(4,6). It is not human subjects research. To identify conditionally independent cell types among all significant cell types implicated we conducted pairwise conditional analyses using MAGMA as previously described (5). GWAS summary statistics were obtained from the most well-powered BPD GWAS available, which includes GWAS meta-analyses of studies with all participants (“combined” n=12,339 cases, n=1,041,717 controls), females only (n=10,025 cases and 547,333 controls), and males only (n=2,260 cases and 485,444 controls).

### Cell type annotations

Consistent with leading approaches to classifying cell types(6,8,9), the cells reported here are transcriptomically defined. Note that neuronal cell type descriptions are either anatomical (i.e., known within the confidence limits of anatomical dissections(6) such as retrosplenial cortex/Broadmann areas 29 and 30) or inferred transcriptomically (e.g., “excitatory”, “layer 2/3”, etc.).

## Results

Ten total cell types were significant after Bonferroni correction: three in the combined analysis, seven in the female analysis, and none in the male analysis. All three Bonferroni-significant cell types in the combined analysis were also significant at FDR *q*<0.05 in the female analysis; of the seven Bonferroni-significant female cell types, two were significant at FDR *q*<0.05 in the combined analysis. All Bonferroni-significant cell types were neuronal, and all were inhibitory (i.e. GABA expressing, versus excitatory/glutamate expressing). Two main categories of neurons were implicated, each encompassing five cell types: LAMP5 interneurons and medium spiny neurons (MSN). Among the LAMP5 interneurons (#287, 270, 271, 288, and 273), three also expressed *LHX6*. The five MSNs (#208, 206, 207, 212, and 213) were primarily of the D2 type (MSN-D2) except for #206, which expressed *DRD1*. As shown in **Figure 1**, the brain regions most enriched for these cell types were in amygdala, cortex, hippocampus, and basal ganglia (nucleus accumbens, caudate, and putamen specifically).

**Figure 1.**
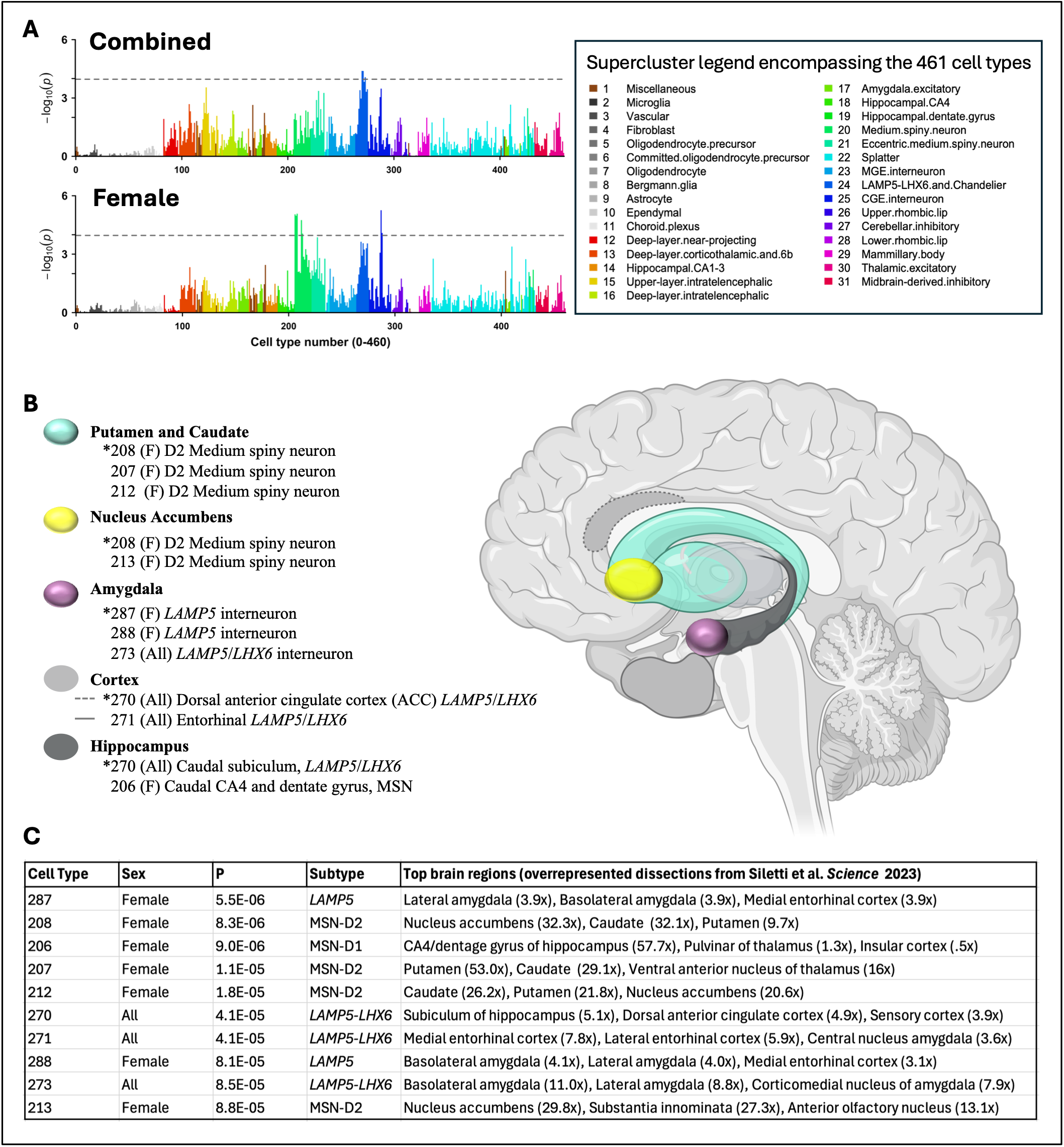
Specific neuronal cell types associated with borderline personality disorder (BPD) **A**. Bar plots depict statistical significance (-log10(*p*)) of the 461 human brain cell types (x-axes) tested in the combined and female analyses. Horizontal dotted lines denote Bonferroni correction (p=.05/461=.0001). Colors denote the 31 superclusters encompassing the 461 “clusters” form Siletti et al. *Science* 2023, which are referred to here as cell types. **B**. Depiction of the top brain regions for each of the ten significant cell types. Within the light-gray shaded regions, dashed outlines indicate the ACC, while solid outlines indicate the entorhinal cortex. Cell types 208 and 270 are depicted twice because their top two regional overrepresentation values were very similar. **C**. Table with summary information about each of the ten significant cell types. *Denotes significant cell types that are relatively independent after conditional analysis.

## Discussion

Consistent with other psychiatric phenotypes, we detected neuronal associations with BPD, and no nonneuronal associations. However, it is very likely that more cell types (neuronal and perhaps non-neuronal) will be found to be associated with BPD using this approach when better powered BPD GWAS become available, as we have shown for successively better powered schizophrenia GWASs(5). Interestingly, the top cellular associations with BPD were not the same top cellular associations that we reported previously for schizophrenia, bipolar disorder, and depression(5). Whereas the top cellular associations for those disorders were also interneurons and MSNs, they were from different interneuron and MSN classes. Regarding interneurons, the top schizophrenia cell types were interneurons from the somatostatin and parvalbumin classes(5), consistent with longstanding reports of altered somatostatin and parvalbumin interneurons in schizophrenia(10). MSN associations were also significant for all these disorders (schizophrenia, bipolar disorder, and depression), but it was D1-MSNs that were most significant. Thus, the prominent associations with LAMP5 interneurons and basal ganglia D2-MSNs stand out for BPD, and suggest that, minimally, these cell types are important for the neurobiology of BPD, and potentially, LAMP5 interneurons and D2-MSNs may be more important for BPD than other major adult psychiatric disorders. Further, while MSNs are classically understood as basal ganglia cell types, we find that cell types annotated as MSNs in the amygdala are more strongly associated with schizophrenia, bipolar disorder, and depression.

LAMP5 (Lysosome-associated membrane protein 5) interneurons are a mammalian-conserved but relatively under-characterized class of predominantly superficial-layer GABAergic neurons that include neurogliaform, single bouquet, canopy, and α7 cells. Neurogliaform cells, which constitute the majority of LAMP5 cortical interneurons, are distinguished by their use of GABAergic volume transmission to generate slow, spatially diffuse inhibition rather than conventional point-to-point synaptic signaling(11). This mode of signaling positions these interneurons as regulators of cortical activity and network state, consistent with recent evidence that they constrain behavioral state-dependent baseline activity in mouse visual cortex (12). Given their laminar localization, LAMP5 interneurons are well positioned to modulate superficial-layer corticoamygdalar projection neurons of prefrontal cortex that govern top-down control of amygdala function (13,14). In borderline personality disorder (BPD), where prefrontal hypoactivity and amygdala hyperactivity are thought to reflect impaired regulatory control (2), one hypothesis is that dysfunction of LAMP5-mediated diffuse inhibitory tone destabilizes cortical gain and degrades the signal fidelity of corticoamygdalar ensembles, thereby weakening prefrontal output and contributing to emotional dysregulation and behavioral impulsivity.

### Analytical notes

Our analysis was not restricted to genome-wide significant signals, rather it leverages information from the millions of SNPs in the BPD GWAS(4). The genetic correlation between male and female subsets was *r*_*g*_=.80 (95% CI *=* 0.53,1.07), suggesting that there might be differences in genetic effects between males and females, though this point estimate was not statistically significantly different from 1. A better powered GWAS of BPD in males will be required to determine if cell types might be different between males and females, but for now the most parsimonious explanation is that the differences in cell types are attributable to differences in statistical power.

### Future directions

Based on our findings with other psychiatric phenotypes, we anticipate more brain cell types will be found to be associated with BPD with better powered GWAS. Importantly, however, we do not anticipate that all or even most brain cell types will be associated; rather, we find that current schizophrenia and depression GWAS appear to be sufficiently well powered, already, to detect the majority of cell types that will be found to be associated (at the currently examined level of cell type granularity), using this method. This gives us greater confidence that the results reported here likely represent the first in a limited set of BPD associated cell types. Next steps include 1) labeling these cell types, *in situ*, in human brain tissue so that relevant receptors and other druggable targets can be characterized, 2) testing the functioning of these cell types *in vivo*, by examining animal homologs and/or *in vitro* using human tissue samples and/or induced neurons, and 3) exploring the potential modulation of these cell types by existing and novel compounds to potentially develop the first, genetics-first therapeutics for BPD.

## Data Availability Statement

All data used and analyzed in this study are publicly available. The snRNA-seq dataset used in this study can be downloaded as described in reference^6^.The GWAS datasets for Borderline Personality Disorder (BPD) is from the Psychiatric Genomics Consortium (https://pgc.unc.edu/for-researchers/download-results/). The code used for our implementation of the MAGMA analysis pipeline is available on GitHub (https://github.com/Integrative-Mental-Health-Lab/linking_cell_types_to_brain_phenotypes).

## Acknowledgements

We thank the participants in the BPD GWAS for their contributions to research, the scientists who established the individual BPD GWAS studies, and the Psychiatric Genomics Consortium (PGC) for sustained pragmatic and scientific leadership in the conduct of GWAS meta-analyses of psychiatric disorders.

## Author information Authors and affiliations

^1^Department of Psychiatry and Behavioral Sciences, Stanford University, Stanford, CA 94305 Hazal Senturk, Hadar Segal-Gavish, Jason Tucciarone, Laramie Duncan

^2^Wu Tsai Neurosciences Institute, Stanford University, Stanford, CA 94305 Laramie Duncan

## Contributions

Planned the paper (LD, HS), wrote and edited the paper (LD, HS, HS-G, JT), conducted analyses and made figures (HS, HS-G, LD), obtained funding (LD).

