## Supplementary text for "Specific basal ganglia, amygdala, and cortical neurons are associated with borderline personality disorder (BPD)"

Cell type annotations

Consistent with leading approaches to classifying cell types, the cells reported here are transcriptomically defined. Note that neuronal cell type descriptions are either anatomical (i.e., known within the confidence limits of anatomical dissections such as retrosplenial cortex, specifically Broadmann areas 29 and 30) or inferred transcriptomically (e.g., “excitatory”, “layer 2/3”, etc.) per annotation pipelines developed by the Linnarsson Lab (<https://github.com/linnarsson-lab/auto-annotation-ah>). Transcriptomically inferred variables are widely used in such studies and are often considered to be factual (e.g. “excitatory” and “inhibitory” neurons inferred based on expression of glutamate and GABA genes, respectively). Additional inferred variables include cell type class (e.g., neuron, oligodendrocyte) and subtype (e.g., “layer 2/3”, “MSN-D1). These are also often considered to be true, but – to varying degrees – need to be validated in human brain tissue.

MAGMA

We employed Multi-marker Analysis of GenoMic Annotation (MAGMA v1.10) software used for gene and gene set analysis of GWAS data(7,15) to complete a two-stage, regression-based procedure to test for associations between BPD, individual genes, and genes used preferentially by human brain cell types. The first stage, gene-level analysis, used a multiple regression approach to calculate gene-level *p*-values that quantify the statistical significance of a gene-phenotype association. The second stage used these gene-level *p*-values to compute *p*-values each cell type by testing for a positive relationship between gene specificity values and gene-level phenotypic associations from the first stage of MAGMA. SNPs were annotated to genes using MAGMA’s single-nucleotide polymorphism (SNP)-gene mapping functionality. Specifically, we employed the same proximity-based criteria used previously(5,15), mapping each SNP to a gene if it resided within a window extending 35 kilobases (kb) upstream of the gene’s transcriptional start site to 10 kb downstream of the end of the gene. MAGMA’s gene property analysis quantified the association strength between a gene and a given phenotype by transforming the *p*-value of a given gene from the gene-level analysis step (𝑃𝑔) into a z-score (𝑍𝑔) using the formula 𝑍𝑔=probit(1−𝑃𝑔). To mitigate potential undue influences of outliers on the analysis, we implemented MAGMA’s default parameters that truncated z-scores 3 standard deviations below and 6 standard deviations above the mean. Gene property analysis was conducted using a linear regression model, expressed as 𝑍=β_0_+𝑃_𝑐_β_1_+𝐶β_2_+𝜀), where 𝑍 represents the aforementioned z-scores of each gene, 𝑃_𝑐_ represents the specificity of each gene in a given cell 𝑐, 𝐶 represents the covariates, and 𝜀 is modeled as a multivariate normal accounting for the LD between genes. Per MAGMA default parameters, specificity values 5 or more standard deviations above the mean were excluded. The covariates in this analysis were gene size, gene density, sample size, inverse mean minor allele count, and their log-transformed values. The final step involved conducting a one-directional test (for positive values) of the coefficient β_1_ to evaluate the association between each of 461 cell types- and a given phenotype. To identify putatively independent signals among all significant results, we conducted pairwise conditional analyses using MAGMA.

Considerations regarding anatomical annotations

Anatomical locations mentioned throughout this paper were obtained from Siletti et al., who sampled the nuclei of 3,369,219 cells across 105 human brain dissections. However, the vast majority of the 461 identified cell types are not exclusive to a single dissection or brain region. Consistent with Siletti et al., dissection annotations were used to quantify enrichment across anatomical locations rather than to definitively localize cell types to single dissections. Donor-level dissection counts were summed to obtain $n_{c,d}$, the number of cells assigned to cell type $c$observed in dissection $d$. We defined $N_{c}$as the total number of cells in cell type $c$ across all dissections, $N_{d}$ as the total number of cells sampled from dissection $d$across all cell types, and $N_{\mathrm{total}}$ as the total number of cells in the dataset. Dissection-level overrepresentation was quantified as ${O/E}_{c,d}=\frac{n_{c,d}/N_{c}}{N_{d}/N_{\mathrm{total}}}$. The top overrepresented dissection was defined as the dissection with the highest ${O/E}_{c,d}$for that cell type and is the dissection highlighted in manuscript figure.
